# Genome-resolved multi-omics insights into top-down microbiome engineering for carbon-upcycling of thermally hydrolysed food waste

**DOI:** 10.1101/2023.05.23.541846

**Authors:** Minxi Jiang, Halil Kurt, Kevin Gilmore, Nathalie A Delherbe, Wendell Khunjar, Lydia Rachbauer, Kartik Chandran

## Abstract

Engineering environmental microbiomes enables carbon upcycling from organic solid waste into valuable products, such as volatile fatty acids (VFAs). This approach leverages the broad metabolic potential of natural communities to process highly heterogeneous substrates. However, solid waste conversion is often constrained by microbial hydrolysis, which limits the breakdown of particulate substrates into soluble compounds available for fermentation. Top-down engineering enhancements, such as thermal hydrolysis pretreatment (THP), can improve feedstock solubility but do not consistently increase total acid yields and may instead shift acid profiles. This highlights our limited understanding of microbiome responses to feedstock-level engineering controls. In this study, we applied genome-resolved multi-omics analysis to link VFA production performance with molecular mechanisms in a top-down engineered microbiome fed with thermally hydrolysed food waste. THP reduced total VFA yield (p = 0.003), accompanied by decreased Shannon diversity (p = 0.03), acid-production potential (DNA; log2FC = −1.3), and acid-production activity (mRNA; log2FC = −0.2). Propionate remained the dominant product with and without THP, consistent with the predominance of the *Prevotella* genus; however, THP selectively enriched two *Prevotella* species with elevated expression of ATP-generating steps in propionate production. THP also increased butyrate and valerate fractions (p < 0.001), together with upregulated reverse β-oxidation for chain elongation driven by CAG-791 sp900320025 and *Megasphaera* sp000417505. <u>Overall, our results provide performance- and molecular-level evidence that THP applied to complex feedstocks didn’t increase total VFA yields, but reshaped microbiome structure and function, maintained propionate dominance, and promoted chain elongation.</u>

## 1. Introduction

The widespread production and inadequate management of anthropogenic solid organic waste including food waste presents a critical environmental challenge, and yet simultaneously offers a vast, underutilized resource for circular bioeconomy (Khanna et al., 2024). Anaerobic fermentation of food waste provides a powerful approach for carbon upcycling by converting organic waste into volatile fatty acids (VFAs), which serve as carbon sources for downstream biological nitrogen removal in wastewater treatment plants and as intermediates for other high-value end-products such as bioplastics and sustainable aviation fuels (Wu et al., 2024; Zhang et al., 2023). This conversion is catalyzed by environmental microbiomes, which contain diverse metabolic capabilities and complex metabolism interactions (Chandran, 2025; Jiang et al., 2024). Compared with single-strain systems or simple synthetic communities, environmental microbiomes offer greater resilience and adaptability for processing real-world wastes with highly heterogeneous compositions.

To optimize microbiome-based fermentation, top-down engineering enhancements can be applied at the system level to steer the substrate and operational conditions experienced by the whole-community (Lyu et al., 2024). For solid waste fermentation, feedstock control is particularly important because microbial hydrolysis is often limiting. Particulate organic matter must first be converted into smaller soluble compounds before being transported into microbial cells and further fermented to volatile fatty acids (VFAs) (Qin et al., 2025; Wilson and Novak, 2009). Thermal hydrolysis pretreatment (THP) is commonly applied to enhance hydrolysis by exposing the feedstock to high temperature and pressure, followed by rapid depressurization (Sahu et al., 2022; Huang and Xiang, 2024; Qin et al., 2025). Although this process increases substrate solubility, its effects on microbiome-driven VFA production remain inconsistent, with reports of both increased and decreased total VFA yields, and variable shifts in acid composition (Castro-Fernandez et al., 2023; Kakar et al., 2025; Penghe et al., 2020; Xiang et al., 2023). These inconsistencies highlight our limited ability to predict microbial community responses to system-level engineering controls, constraining the scalability of microbiome-engineering-based approaches for solid waste upcycling. Advancing this field requires a deeper mechanistic understanding of dynamic shifts in microbiome structure, function and metabolic pathways after the inclusion of system-level pretreatment of feedstocks (Atasoy et al., 2024).

Multi-omics approaches offer powerful insights into microbiome composition and function, yet their application in solid-waste carbon-upcycling has concentrated on biogas rather than VFA production (Akinsola et al., 2025; Liang et al., 2020; Xia et al., 2018; Zhang et al., 2021). Limited research has evaluated the VFA-producing microbiomes fed with substrate after different pretreatments, including microaeration (Amin et al., 2024), ferrate and alkaline pretreatment (Li et al., 2018), enzymatic pretreatment (Zhang et al., 2025), and THP. In THP studies, enrichments of diverse acidogenic bacteria are reported such as Firmicutes, particularly Clostridia (e.g., *Clostridium sensu stricto*, *Caproiciproducens*, *Paraclostridium*, and *Sedimentibacter*) (Castro-Fernandez et al., 2023; Shi et al., 2022; Wan et al., 2020), as well as Coprothermobacteria and Thermotogae (Kakar et al., 2025). These findings remain descriptive, relying on DNA-derived taxonomy, functional predictions from off-the-shelf bioinformatic tools (Castro-Fernandez et al., 2023), or statistical correlations between produced products and enriched taxa. However, in mixed communities fed complex organic substrates, gene abundance does not necessarily reflect expression activity, and population shifts can be decoupled from targeted metabolic functions (Abreu and Taga, 2016; Jiang et al., 2024). Consequently, accurate assignment of functionally active organisms requires transcription-level or other multi-omics validation of genes, enzymes, and pathways, which remain limited in current literature (Boehm et al., 2024; Chandran, 2025).

In this study, we applied integrated metagenomics, metatranscriptomics and 16S rRNA sequencing approaches to comprehensively characterize genome-resolved functional shifts in top-down engineered acidification microbiomes fed with THP-treated food waste. We hypothesized that changes in activity of a specific acid production pathway, rather than relative abundance alone, would better explain VFA yields and speciation. We operated two mesophilic reactors (target operating temperature of 37 ℃) at a hydraulic retention time (HRT) of 2 days to suppress methanogenesis: one fed with THP-treated food waste and the other with untreated food waste (THP and non-THP reactors, respectively). Feedstock characteristics and VFA production performance were monitored. Microbial community dynamics were assessed by 16S rRNA gene sequencing for structural diversity, metagenomics for reconstructing genome-resolved MAGs, and metatranscriptomics for quantifying active gene expression through transcript mapping to assembled MAGs. By comparing acidification networks before and after THP, this study provides insights into key species, essential genes and pathways that drive VFA production from THP-treated feedstocks.

## 2. Materials and Methods

### 2.1 Food waste sampling and THP conditions

Three batches of food waste (FW1, FW2, and FW3) were collected from the University café and homogenized with a kitchen blender on the day of collection. THP was performed the following day at Bucknell University (Lewisburg, PA) using a pilot-scale THP unit. Each batch was subjected to steam pressurization at 6 bar (around 165 ℃) for 40 minutes, with manual venting every 10 minutes. This was followed by a flash step with rapid pressure release, after which the vessel contents were expelled into a container. Both THP- and non-THP treated food waste were aliquoted into 250-mL plastic bottles and shipped overnight on dry ice to Columbia University. All food waste was stored at –20 ℃ to prevent spoilage until use. All batches underwent the same freeze-thaw cycles prior to characterization and feeding to ensure consistency.

### 2.2 Reactor set-up and operations

Two 6-L semi-continuous reactors were operated at 37.0 ± 2.0 ℃ for 38 days, and pH was maintained at 6.20 ± 0.05 using a feedback-controlled monitoring system with 1M NaHCO_3_ buffer. Three food waste batches were fed sequentially to both reactors, with each batch split into THP-treated and non-THP-treated feed for paired reactor operation (**SI Methods**). To minimize confounding effects from feedstock variability, the organic loading rate (OLR) was maintained consistently at 6.25g total COD/L/d between the two reactors across all three food waste batches, which is within a typical range reported for fermentation reactors (Magdalena et al., 2019). Both reactors were operated at a hydraulic retention time (HRT) of 2 days. Because the reactors were continuously mixed and no solids-liquid separation was performed before effluent withdrawal, the solids retention time (SRT) was equal to the hydraulic retention time (HRT). Details of the automated pumping system used to achieve the targeted HRT are provided in the **SI Methods**. This short retention time was maintained to limit methanogen growth (De Groof et al., 2021). All reactors were inoculated with 500 mL effluent from a previously operated mixed-culture arrested anaerobic digestion reactor (Jiang et al., 2024).

### 2.3 Performance analysis

The characteristics of THP and non-THP treated food waste were measured for each batch following previously reported methods (Jiang et al., 2024), including chemical oxygen demand (COD), total solids (TS), volatile solids (VS), carbohydrates, proteins, lipids, total ammonia nitrogen (TAN), and total carboxylic acids (tCA) (**Table S1 and SI Methods**). Effluent metabolites were monitored two to three times per week, including soluble COD (sCOD), volatile fatty acids (VFAs), and methane levels in the headspace (**SI Methods**). Bioreactor performance was assessed based on total VFA conversion efficiency, methane yield, VFA accumulation level, and individual acid proportions (**SI Methods**). All measurements for both influent and effluent were conducted in duplicate.

### 2.4 16S rRNA gene amplicon, metagenomics and metatranscriptomics sequencing

Biomass was collected at three time points (days 17, 23, and 31) after stable VFA production was achieved, as indicated by effluent VFA concentrations showing less than 8% relative standard deviation among samples collected within a week (**Figure 2**). Biomass from 2-mL samples were pelleted by centrifugation at 13000 xg, and stored at −80 ℃ with RNAprotect (Qiagen, MD, USA). DNA and total RNA were separately extracted using the DNeasy PowerSoil kit (Qiagen, MD, USA) and RNA PowerSoil Kit (MO BIO laboratories, Carlsbad, CA), respectively. Nucleic acids were quantified with Qubit 1X dsDNA broad-range assay kit and Qubit RNA HS Assay (Fisher Scientific). For 16S rRNA gene amplicon sequencing, the primer set 515F/806R targeting the V4 region of both bacteria and archaea was used to amplify the 16S rRNA genes (Caporaso et al., 2011), and sequencing was performed on an Ion Torrent PGM (Thermo Fisher, MA, USA). For metagenomic and metatranscriptomic sequencing, biomass from the three time points was pooled in equal mass to obtain a final input of 500 ng of DNA and RNA per feedstock condition (THP and non-THP). The quality of the pooled DNA and RNA was assessed using a bioanalyzer (Agilent 2100). Library preparation and sequencing was performed at GeneWiz (Azenta Life Sciences), on the Illumina Hiseq platform with 2 x 150bp read lengths.

### 2.5 Bioinformatics analysis

For 16S rRNA gene sequencing, data were processed using QIIME2 (version 2019.10). OTUs were identified at a 97% similarity. Taxonomic classification was performed with the Silva.132 database, and results were visualized in R (v3.6.3) or Python (v3.8.0) with custom scripts.

For metagenomics analysis, raw paired-end reads were quality filtered and adapter-trimmed using Trimmomatic v0.39 (mean quality ≥ 15, minimum read length ≥ 36 bp). Read-based functional analyses were performed by aligning high-quality reads with DIAMOND (v2.0.14), and functional annotations were generated in MEGAN using the eggNOG database (Huerta-Cepas et al., 2019) following a previously published protocol (Jiang et al., 2024). High-quality reads were also assembled into contigs using both MEGAHIT v1.2.9 and SPAdes v3.13.1, retaining contigs > 1000 bp. Assembly quality was assessed with Metaquast (v5.2.0), and coverage depth of contigs were calculated. Co-binning of THP and non-THP sequences was performed with metaWRAP v1.3.2 (binning module using MetaBAT2 v2.9.1, CONCOCT v0.4.0, and MaxBin2 v2.2.4), followed by metaWRAP bin_refinement to consolidate bins. The generated metagenome-assembled genomes (MAGs) quality were assessed with CheckM (v1.0.7), and bins with ≥85% completeness and ≤5% contamination were retained (classified here as medium-to-high quality following MIMAG criteria) (Bowers et al., 2017). Redundant MAGs were dereplicated at 95% average nucleotide identity using dRep (v3.5.0), resulting in final 21 MAGs (assembly and QC statistics in **Table S2**). Taxonomy was assigned to each MAG using GTDB-Tk (v2.1.1). Genes were predicted using Prodigal (v2.6.3) (Hyatt et al., 2010) in metagenomic mode, and translated amino acid sequences were annotated by Prokka (v1.14.6) and KofamScan (v1.3.0) against the KEGG database. Additional pathway summaries were generated with METABOLIC-G (v4.0), mapping KOs/ECs to KEGG modules and carbon-metabolism pathways (**Table S3**).

RNA-seq reads were similarly quality filtered and rRNA sequences were removed with SortMeRNA (v4.3.7). Expression was quantified by mapping reads to MAG with Bowtie2 (2.5.4) and counted with customized python script to generate TPM per gene (**SI Methods**).

Seven assembled MAGs classified as *Prevotella* species were analyzed for functional genomic and synteny comparison using Anvi’o (v8) (Eren et al., 2015). Coding sequences (CDSs) were identified by Prodigal (v2.6.3), and functional annotation was assigned based on NCBI’s Clusters of Orthologous Groups (COG) (Galperin et al., 2021) and KEGG KOfam KOALA (Aramaki et al., 2020). Orthologous groups (OG) were determined through all-vs-all amino acid homology comparisons using a minbit of 0.5 and the MCL inflation of 5, optimized for within-genus comparison. Core versus unique OGs were distinguished among the MAGs. The unique OG subsets were further inspected for functional enrichment based on KOfam annotations. Additionally, the whole genome similarity of seven MAGs was assessed using anvi-compute-genome-similarity, utilizing the PyANI similarity metric to calculate the average nucleotide identity (ANI), ANI alignment coverage and ANI full percentage (ANI x alignment coverage) (Pritchard et al., 2015).

To specifically investigate volatile fatty acid (VFA) production, a customized acidification metabolic pathway database was compiled from KEGG and relevant literature (**Table S4**). KOs associated with the acidification process were identified across all MAGs, the expressed pathways were reconstructed and visualized in Adobe Illustrator (v2025). Reconstruction was focused on pathways beginning with pyruvate leading to individual VFA, as pyruvate is a central intermediate in food waste fermentation across diverse organic fractions. The scope was also primarily placed on carbohydrate metabolism, while protein degradation was simplified to include only terminal reactions contributing to VFA production after proteins are broken down into simpler carbon structures.

### 2.6 Statistical analysis and data availability

Statistical differences between THP and non-THP groups were evaluated using Welch’s t-test. If normality was not met, the Mann–Whitney U test was applied instead. All statistical tests were performed using the Python package scipy.stats, and significance was determined at a 95% confidence interval (p < 0.05, **Table S5**). All raw sequencing data were deposited in the NCBI Sequence Read Archive (SRA) under accession number SRS17432456. The analysis workflow is available at https://github.com/mj2770/Multiomics_MMCs_fermentation.

## 3. Results and Discussion

### 3.1 Changes in food waste characteristics after THP

The influent total COD concentrations for both the THP and non-THP reactors were kept consistent to maintain the target OLR across different FW batches (**Figure 1**, COD). However, as expected, THP consistently increased feedstock solubility as evidenced by higher soluble COD and decreased total solids (TS) (p < 0.05, **Table S5**), consistent with previous research (Castro-Fernandez et al., 2023; Huang and Xiang, 2024). Varying degrees of organics degradation were also observed across different FW batches. Soluble carbohydrates concentration increased in all three batches, while the soluble protein increased only in FW1 after THP. Meanwhile, the total ammonia nitrogen (TAN) concentration increased in FW1 and FW2 (p < 0.05, **Table S5**), which indicated potential protein degradation during THP (Shi et al., 2022). The measured TAN levels are at the lower end of values reported for food and kitchen waste (< 60 to > 5000 mg NH₄⁺-N/L) (Ebel et al., 2025), likely reflecting specific composition properties of our collected feedstock. Lipid content (% wet weight) decreased in FW1 and FW2 after THP (p < 0.05, **Table S5**), alongside higher total carboxylic acid (tCA) levels (**Figure 1**), suggesting possible lipid conversion to carboxylic acids during THP (Yu et al., 2016). Notably, both the free ammonia (< 1.24 mg FA-N/L, calculated from < 500 mg NH₄⁺-N/L at pH 6.2 and 37°C (Liu et al., 2019)) and total carboxylic acid (< 0.4 g COD-tCA/L) in THP-treated feedstock were below reported inhibitory thresholds for microbial activities during VFA production process (Shi et al., 2017). However, these compounds may continue to accumulate during subsequent fermentation, which was not assessed in this study. Overall, although the three batches were operated at similar organic loading rates, their characteristics differed markedly before and after THP (**Figure 1**, PCA plot). This underscores the high variability in real food waste and the resulting impact on THP performance itself (Chen et al., 2018).

**Figure 1.**
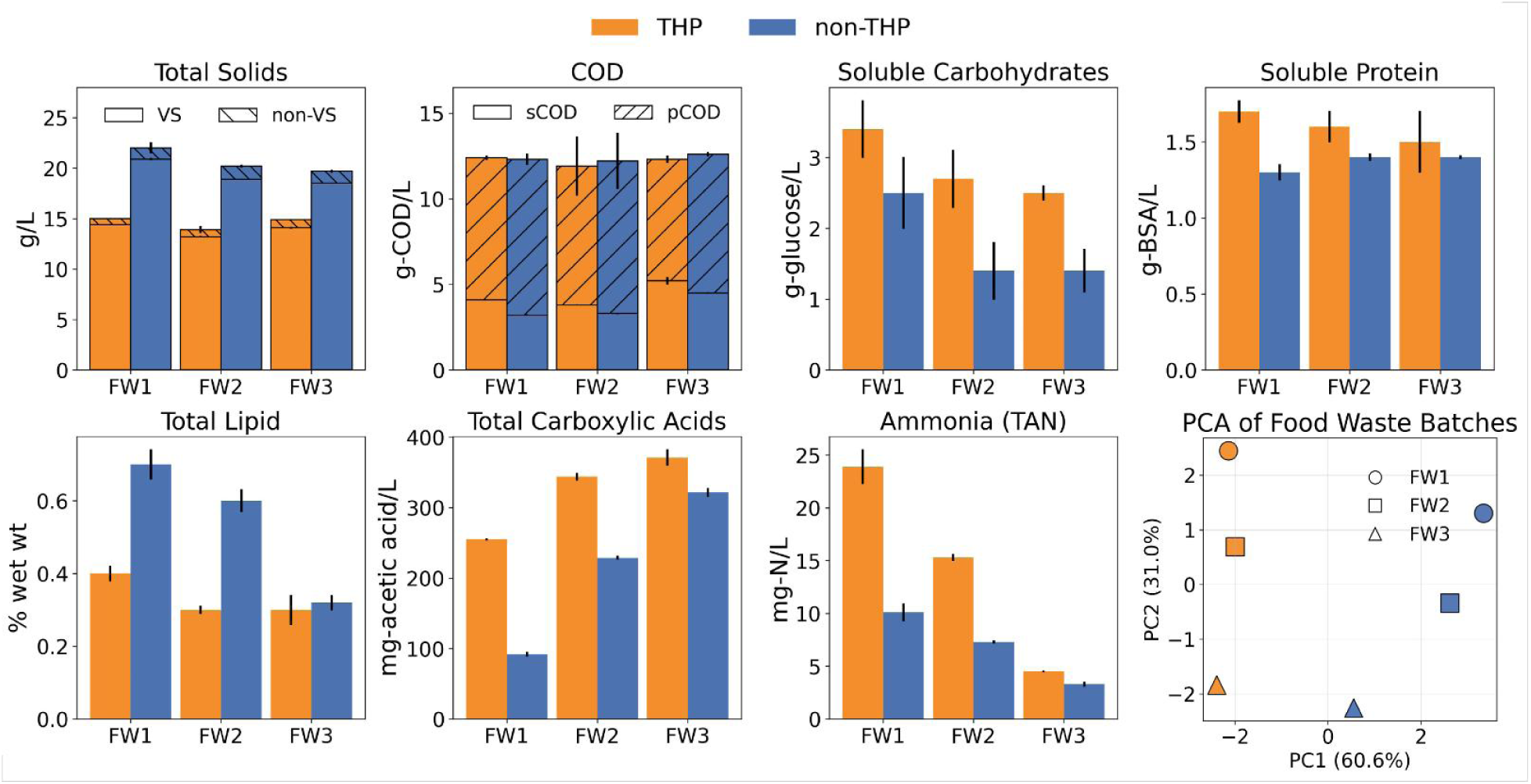
Effects of THP on food-waste composition across three batches. Colors indicate treatment: THP (orange) and non-THP (blue). **Top row**, left to right: total solids (TS) partitioned into volatile solids (VS, open) and non-VS (hatched); chemical oxygen demand (COD) partitioned into soluble COD (sCOD, open) and particulate COD (pCOD, hatched); soluble carbohydrates; soluble protein. **Bottom row**, left to right: total lipid; total carboxylic acids (tCA; as acetic acid equivalents); total ammonia nitrogen (TAN); PCA of standardized food waste property metrics (symbols denote FW1-FW3). Error bars represent standard deviations from duplicate measurements.

### 3.2 VFA production performance

Although food waste batches varied in characteristics, when fed at a constant total COD, both THP- and non-THP-fed reactors reached stable VFA production after 15 days, with effluent VFA concentrations varying by less than 8% (**Figure 2a, 2b**). During this period, the THP reactor showed a significantly lower average effluent VFA concentration than the non-THP reactor (4.8 ± 0.4 vs. 6.1 ± 0.9 gCOD/L, respectively; n = 8, p = 0.004; **Table S6**). Because both 6-L reactors were operated at the same organic loading rate, this lower concentration indicates reduced total VFA conversion efficiency in the THP reactor, decreasing from 48.8 ± 6.8% to 38.6 ± 3.2% (p = 0.003; **Table S6**). VFA accumulation level, expressed as the ratio of VFA concentration to effluent soluble COD, was also lower in the THP reactor than in the non-THP reactor (67.2 ± 5.8% vs. 85.1 ± 9.2%, respectively; Figure 2b and Table S6). Similar reductions in VFA yield after THP have been reported for sewage sludge, suggesting that a substantial fraction of solubilized organics may be converted into compounds other than VFAs (Castro-Fernandez et al., 2023). Although other soluble products were not quantified in this study, alcohol accumulation has been reported during fermentation of hydrothermally pretreated cellulose and sludge (Zhang et al., 2023; Zhou et al., 2024). Additionally, recalcitrant organics could also be formed after THP, such as protein-derived soluble nitrogen compounds, which persist in the feedstock and are poorly converted to VFAs (Zhang et al., 2020).

**Figure 2.**
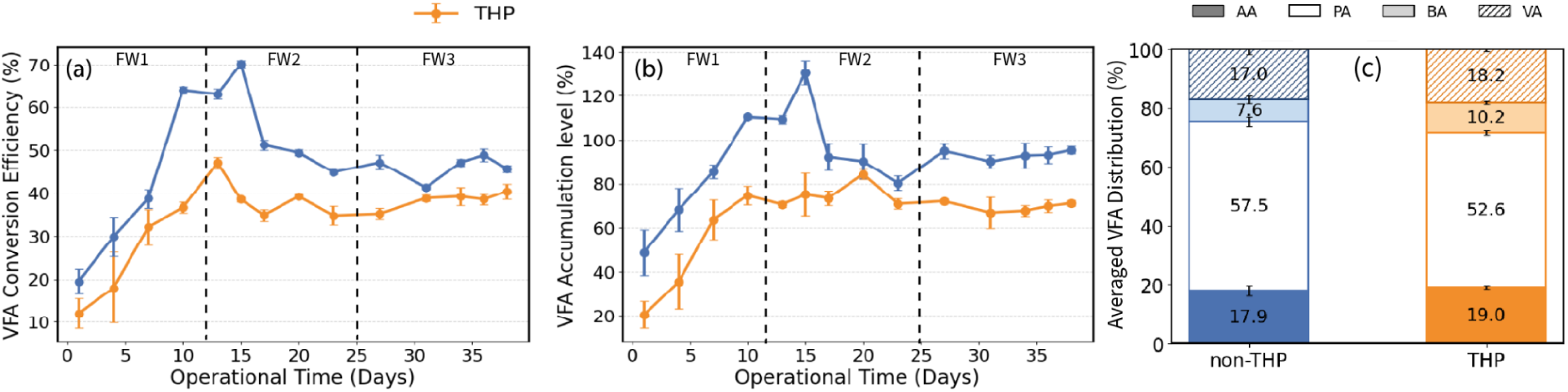
VFA production performance and composition in THP and non-THP reactors during food waste fermentation. **(a)** VFA conversion efficiency (%) and **(b)** VFA accumulation level (%) over operational time (38 days) in THP (orange) and non-THP (blue) reactors. Error bars represent standard deviations from duplicate measurements. VFA accumulation level (%) higher than 1 mg VFA/mg sCOD reflect differences between sCOD and IC-based VFA measurements, which rely on theoretical COD equivalents for each acid (**SI Methods**). **(c)** Acid composition in produced VFAs shown as the relative distributions % (**SI Methods**) of acetic acid (AA), propionic acid (PA), butyric acid (BA), and valeric acid (VA). Values are averaged between 15 - 38 days after reactors reach steady state.

In terms of acid composition, acetate (AA), propionate (PA), butyrate (BA), and valerate (VA) were detected in both reactors, with PA remaining the dominant product regardless of THP (non-THP: 57.5 ± 1.8%; THP: 52.6 ± 0.9%; n = 8, **Figure 2c and Table S6**). However, the fraction of PA (C3) in produced total VFAs was significantly lower in THP reactor than in non-THP reactor, whereas the BA (C4) and VA (C5) fractions were higher (BA: non-THP 7.6 ± 1.3% and THP 10.2 ± 0.5%; VA: non-THP 17.0 ± 1.8% and THP 18.2 ± 0.6%) (n = 8, **Figure 2c and Table S6**). Although the magnitude of this shift was modest, it still suggests that a greater fraction of carbon flux was directed toward more elongated products (> C4) than short-chain acids (C2 and C3) in the THP reactor. The following multi-omics results also showed increased expression of genes involved in reverse β-oxidation pathways, although mainly from two low-abundance organisms (**Section 3.6**). Similar enhancement of chain elongation after thermal hydrolysis pretreatment have also been reported in sludge systems, where *in situ* produced ethanol after THP significantly enhanced C6-C8 carboxylic acid production without external electron donor addition, and PICRUSt-based functional prediction indicated increased activity of the reverse β-oxidation pathway (Zhang et al., 2023).

Lastly, no methane production was detected in either reactor (**Table S6**), likely because the 2-day HRT was insufficient to support methanogen growth. This observation aligns with prior findings that methanogens are more susceptible to wash-out under short HRTs, especially in acidogenic or fermentative systems with suboptimal pH (Gaby et al., 2017). The headspace hydrogen gas was not directly measured in this study. However, as hydrogen partial pressure is essential in maintaining VFA formation (Welsh et al., 2025) and potential chain-elongation in fermentation communities (Baleeiro et al., 2021), we assessed the hydrogen production and consumption pathways, as reported in the following multi-omics analysis (**section 3.5**).

### 3.3 The impact of THP feedstock on microbial ecology of anaerobic fermentation processes

The 16S rRNA gene sequencing results included three time points collected between operational days 17 and 31, when reactors reached relatively stable performance (**Figure S1**). Reactors receiving THP feedstock showed a significantly reduced Shannon diversity compared to the non-THP reactor (n = 3, p = 0.03, **Figure S1**), which likely reflects the loss of indigenous microbes in food waste during the THP sterilization (Shi et al., 2022). Although *Prevotella* genus dominated both reactors, its abundance was significantly higher in the THP reactor than in the non-THP reactor (80.32 ± 3.32% v.s. 54.91 ± 7.52%; n = 3, p < 0.05, **Figure S1**). The THP reactor also showed higher *Mitsuokella*, whereas non-THP reactor was enriched in uncultured members of the family Succinivibrionaceae and in the genus *Bacteroides* (n = 3, p < 0.05, **Figure S1**). Methanogens were not detected in either reactor, consistent with effective washout under the 2-day HRT. We co-assembled the two metagenomic results (as described in Methods) from the THP and non-THP reactors and recovered 21 good quality MAGs (**Figure 3**). Read back-mapping to these MAGs mirrored the 16S-based abundance differences between reactor types while providing species-level resolution. In the non-THP reactor, *Succinivibrio hippei* and *Bacteroides* sp007896885 were enriched relative to the THP reactor. *Prevotella* genus were prevalent in both reactors, yet individual species were differentially enriched between reactor types. *Prevotella* sp002305235 and *Prevotella* unclassified 2 were highly enriched in THP, whereas *Prevotella* sp018054505, *Prevotella* unclassified 1, and *Prevotella* unclassified 3 predominated in non-THP (**Figure 3**). These results suggest niche differentiation among *Prevotella* lineages.

**Figure 3.**
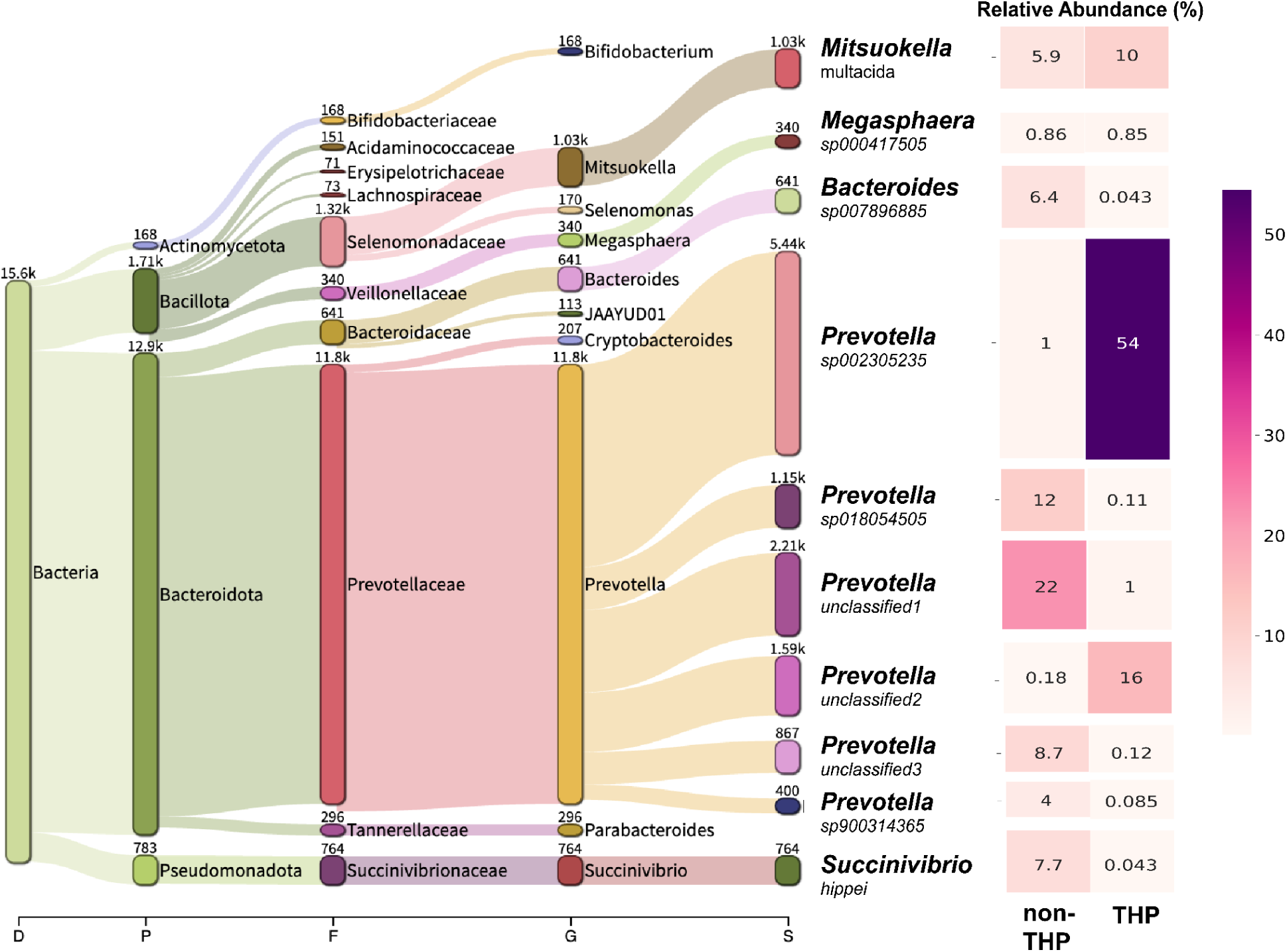
Species-level taxonomic profiles of MAGs in THP and non-THP reactors operated at an HRT of 2 days. Relative abundances of Top 10 species are shown, others are only listed on genus level. The Relative abundance presented here is the proportion of mean coverage of each MAG calculated by coverM (v0.7.0).

### 3.4 Community-level activities associated with specific acidification metabolic networks and essential cellular functions

The community-level activities were first compared using eggNOG functional categories (Huerta-Cepas et al., 2019). Four major functions exhibited higher transcript levels in the THP reactor compared to the non-THP reactor, including *inorganic ion transport and metabolism*, *energy production and conversion*, *secondary metabolite biosynthesis*, and *post-translational modification* (**Figure S2**; mRNA TPM log₂FC > 0.5 and each category representing >10% of total mRNA TPM). However, these categories are too broad to directly link with VFA production, highlighting the need for targeted analyses using a defined acidification metabolic network (**Table S4 and Figure 4a**). Within this network, summed DNA and mRNA TPM values of all genes were lower in THP reactors, while the mRNA/DNA ratio was higher (**Table S7**, log₂FC = –1.3, –0.2, and 1.1, respectively). This suggests that although the overall genomic potential and activity was lower in the THP-associated reactor compared to control, the remaining microorganisms exhibited elevated per-genome transcriptional level of acidification.

**Figure 4.**
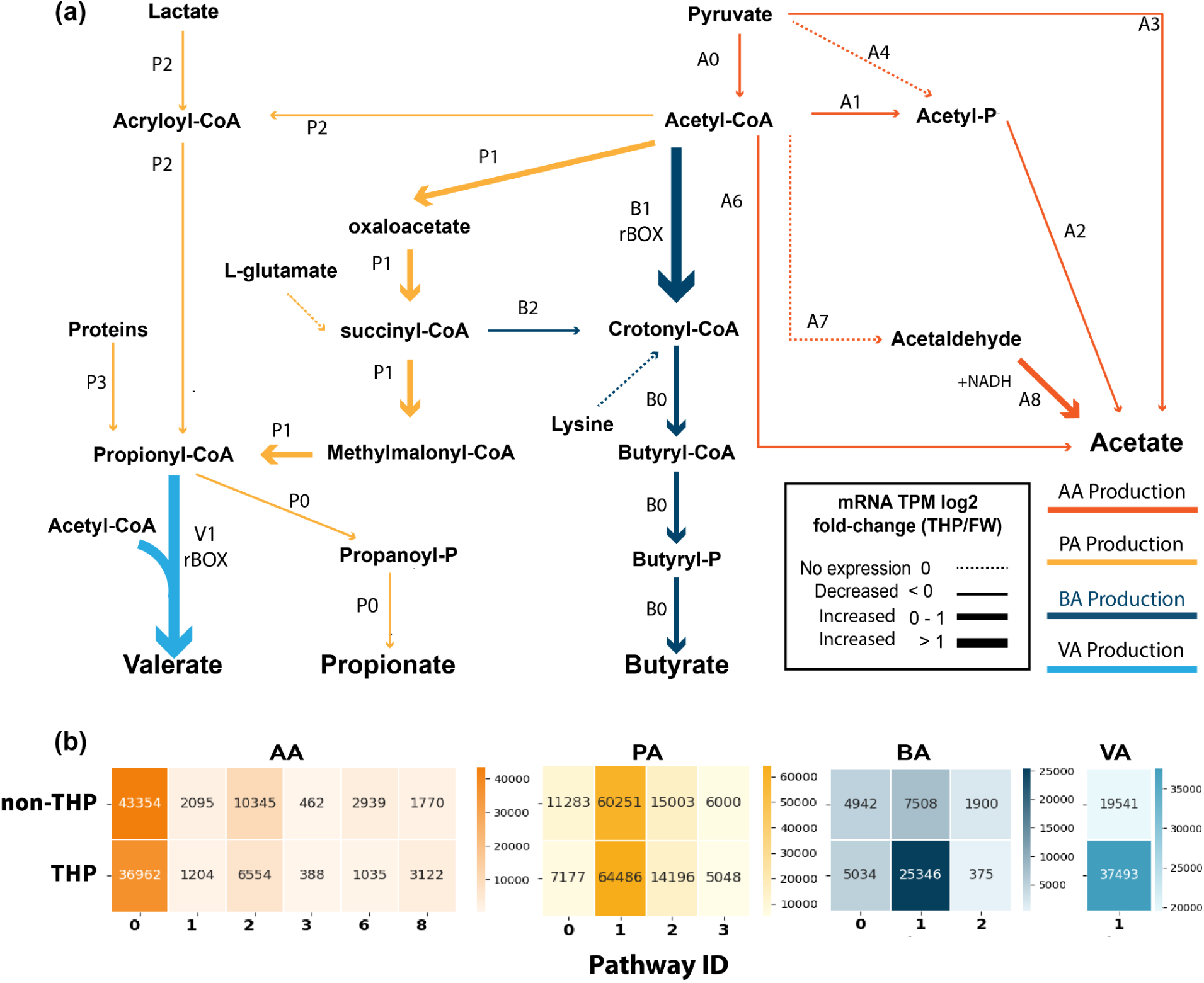
Acidification metabolic network and the changes of community-level transcription profile after THP. **(a)** Schematic of the acidification metabolic network, with pathway activity fold change represented by the thickness of arrow lines (thicker lines indicate increased fold change in mRNA gene expression). Dashed arrows indicate pathways with no detected mRNA expression. Pathways are color-coded based on their end-products. **(b)** Heatmap showing the summed mRNA TPM for each individual pathway involved in acid production (detailed pathway information is defined in **Table S4**). Annotated numbers in each cell represent the summed mRNA TPM of genes involved in each defined pathway.

Although the main routes for the production of individual VFA were conserved between THP and non-THP reactors (**Figure 4b**), their expression levels shifted in the THP reactor (**Figure 4a**; line thickness indicates mRNA TPM fold change after THP). In both reactors, acetate was primarily produced from pyruvate conversion to acetyl-CoA (A0), followed by the acetyl-phosphate to acetate route (A2, **Figure 4a**). Most acetate pathways were downregulated, except acetaldehyde to acetate, which was upregulated in the THP reactor (**Figure 4b**, log2FC = 0.8). We also observed a slight upregulation of the ethanol-to-acetaldehyde step (**Figure 4b**, log₂FC = 0.10), which could potentially provide additional NADH via ADH (alcohol dehydrogenases) and the subsequent ALDH-derived (aldehyde dehydrogenase) reaction for reverse β-oxidation (rBOX) and other NADH-dependent processes in the THP reactor (Sun et al., 2024). The D-lactate to pyruvate pathway was also modestly upregulated (**Figure 4b**, log₂FC = 0.2), indicating lactate as another potential electron-donor to fuel the rBOX pathways (Wang et al., 2022). Propionate was primarily produced through the methylmalonyl-CoA/succinate (MMC) pathway (P1) in both reactors, consistent with prior reported pathways of propionate production from enzyme-pretreated food waste fermentation (Zhang et al., 2025). The summed mRNA TPM involved in this pathway was slightly upregulated after THP, while the final step converting propionyl-CoA to propionate was downregulated in the THP reactor (P0; log₂FC = – 0.65; **Figure 4b**). Butyrate and valerate were produced mainly via the reverse β-oxidation pathway, which was strongly upregulated in the THP reactor (B1 and V1; **Figure 4b**, log₂FC > 1.7; **Figure 4a**). This molecular-level evidence is consistent with the observed increase in butyrate and valerate fractions following THP treatment. Alternative routes for butyrate and valerate production such as lysine degradation or succinyl-CoA conversion were minimally detected. Acyl carrier protein (ACP)-based fatty acid synthesis pathways were not examined, as they commonly represent canonical long-chain fatty acid biosynthesis for membrane lipids rather than fermentation products (Chan and Vogel, 2010). Overall, the community-level mRNA TPM patterns within our custom acidification networks could suggest which pathways were more active in the production of specific acids. Shifts in mRNA TPM before and after THP, both across the full network and within individual pathways, are also partly consistent with the observed performance changes of reduced total acid yields and elevated BA and VA fractions. However, these associations remain tentative, because a more robust interpretation would require metabolic modeling that captures the full functional repertoire of each community member from good quality MAGs, which is challenging for mixed-culture microbiomes due to the limited resolution of DNA- and RNA-sequencing.

### 3.5 Genome-resolved insights into dominant propionate production and selective enrichment of *Prevotella* species

In addition to the community-level analysis of acidification, we further examined individual organism-level activity, with a particular focus on MAGs contributing to the dominant production of propionate in both THP and non-THP reactors. As described in Section 3.4, propionate was mainly produced through the methylmalonyl-CoA (MMC) pathway in both reactors (P1, **Figures 4a**). Mapping of MMC-pathway genes indicated that the required enzymes were broadly encoded and expressed across the community, with different MAGs covering complementary steps of the pathway. The clustering of gene expression based on presence-absence patterns revealed four distinct groups of MAGs with incomplete or divergent propionate-producing capacities (**Figure S5**), supporting a distributed, community-level basis for MMC-mediated propionate formation.

Notably, *Prevotella* genus remained the dominant genus in both THP and non-THP reactors, consistent with the propionate-dominated VFA profiles. Previous studies in food waste fermentation and human gut systems have also implicated *Prevotella* as a major contributor to propionate production in *Prevotella-*dominated microbiome, especially under carbohydrate-rich feedstock (Adamberg and Adamberg, 2024; Chen et al., 2017). Accordingly, we analyzed the mRNA profile of seven assembled *Prevotella* MAGs to evaluate their specific contribution to propionate production (**Figure 5**). We observed that different *Prevotella* MAGs expressed part of the complete MMC pathway, and 6 of the 7 assembled bins actively expressed gene *ackA*, which is involved in producing propionate from propionyl phosphate (propionyl-P) (**Figure 5, heatmap**). THP deactivated this step in bins 11 and 12. Notably, two *Prevotella* MAGs (bins 17 and 21; *Prevotella* sp002305235 and *Prevotella* unclassified 2) exhibited a distinct transcriptional pattern within the MMC pathways, with reduced expression of *frdA/B/C* alongside upregulation of *sucC* and *sucD* (**Figure 5, red highlight**). These two upregulated genes encode enzymes that catalyze the conversion of succinate to succinyl-CoA and generate ATP via substrate-level phosphorylation (Schürmann et al., 2011). Notably, these same two *Prevotella* MAGs were also the ones most strongly enriched after THP treatment (**Figure 3**). Considering that succinate is an essential cross-feeding intermediate in many fermentation communities (Wei et al., 2023), we hypothesized that this transcriptional shift may confer a growth advantage by bypassing the fumarate-reduction step and instead relying on exogenous succinate supplied by other community members, channeling it directly into succinyl-CoA for ATP production.

**Figure 5.**
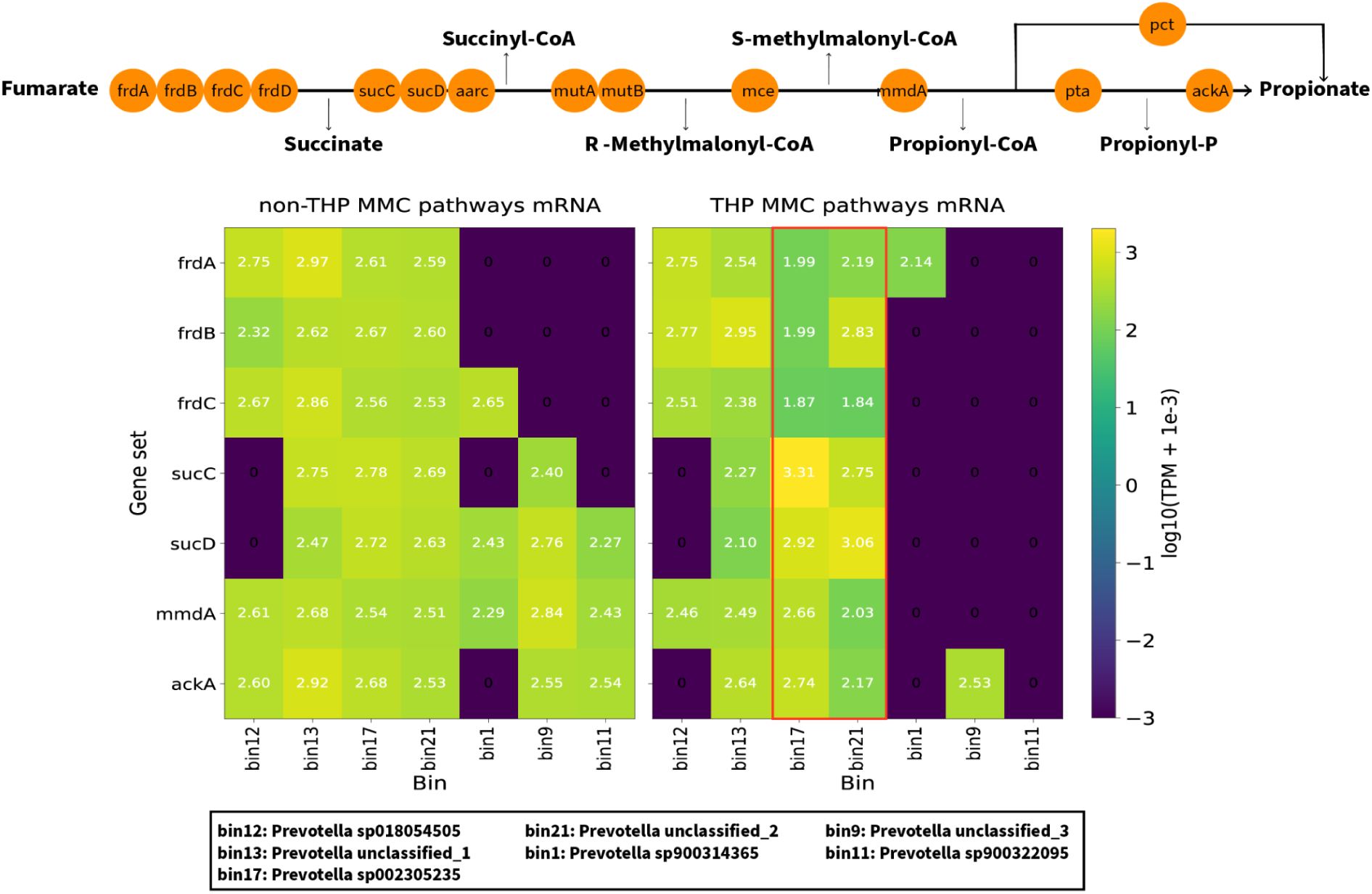
Genome-resolved propionate production transcriptional activity of *Prevotella* MAGs. Expression profiles of key metabolic genes in the propionic acid production through the MMC pathway (succinate→propionate) across *Prevotella* bins under non-THP and THP conditions. The schematic (top) illustrates the fumarate-to-propionate conversion pathway. The heatmaps (bottom) show log10-transformed TPM values of pathway genes across seven *Prevotella* MAGs (bins 12, 13, 17, 21, 1, 9, 11). MAG taxonomy identities are listed below the heatmap.

To further investigate why only these two *Prevotella* species (bin 17 and bin 21) were selectively enriched after feeding with THP food waste, we also performed a DNA-level pangenomic analysis. These two species shared the highest genome-wide ANI (75%; **Figure S6**) relative to the other *Prevotella* species. Examination of their unique gene sets revealed 33 shared KOfam-defined functions, including prominent starch-binding outer membrane proteins from the SusD/RagB family (K21572) and a TonB-dependent starch-binding protein (SusC, K21573) (**heatmap in Figure S6**). These features may provide an additional growth advantage under THP by enhancing starch binding and uptake, however, the precise functional validation of these genes remain to be investigated.

### 3.6 Genome-resolved understanding of niche partitioning for chain elongation

To identify the organisms responsible for the elevated BA and VA fractions and the upregulation of reverse β-oxidation (rBOX) pathways after THP, we examined chain-elongation-related functional modules in the assembled MAGs (**Figure 6a**). These modules included the reverse β-oxidation (rBOX) pathways, electron-donor metabolism (e.g., H_2_-associated electron transfer and redox balancing through Ech complex, lactate and ethanol oxidation to acetyl-CoA and NADH), and energy-conservation systems (Rnf complexes) in the assembled MAGs (**Figure 6a**) (Baleeiro et al., 2023; Scarborough et al., 2020; Wang and Yin, 2022). Only genes expressed in at least one MAG were retained for further comparison analysis. The full bin-wide expression profiles of rBOX-related functional modules are provided in **Figure S3**.

**Figure 6.**
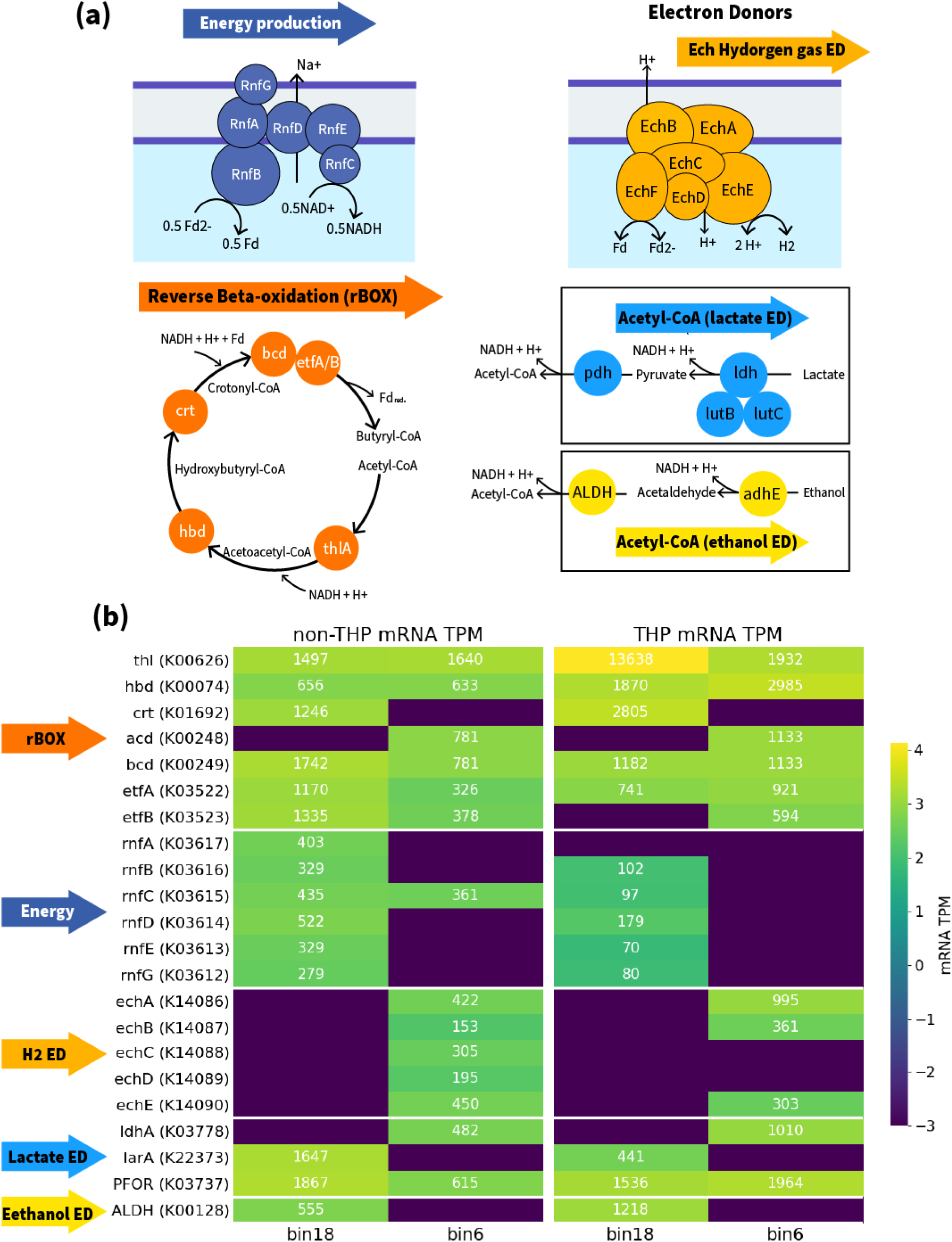
Functional modules, genomic context, and transcriptional profiles of reverse β-oxidation under THP and non-THP conditions. **(a)** Schematic of defined functional modules required to complete reverse β-oxidation (rBOX) and associated supporting pathways, including electron-donor metabolism (hydrogen gas, lactate, and ethanol oxidation) and energy-conservation complexes (Rnf). **(b)** Heatmaps of mRNA expression (TPM, log_10_ scale) for selected genes in bins 18 and 6, under non-THP and THP conditions. Rows are grouped by pathway (rBOX, Rnf, H_2_ donor through Ech, lactate donor, and ethanol donor). Only genes with non-zero expression in the plotted bins are shown; full bin-wide expression data are provided in

Among all MAGs, only *Megasphaera* sp000417505 (bin 18) and CAG-791 sp900320025 (bin 6, under Lachnospiraceae family) encoded and expressed nearly complete rBOX pathways (**Figure S4**), making them the most likely chain elongators in our reactors. In *Megasphaera*, the full rBOX gene set was recovered but dispersed across contigs, with only *hbd-crt* co-localized (**Figure S4**). As this genus has been widely reported to perform chain-elongation through the rBOX pathways (Jeon et al., 2025), this fragmentation of this co-localized rBOX gene set might reflect assembly limitations due to low abundance of this species. All rBOX genes were expressed in *Megasphaera*, with *thlA* upregulated nearly 10-fold in THP reactors (**Figure 6b**), which corresponded to the elevated butyrate and valerate fractions in the product profile. Among the pathways that could supply reducing equivalents for rBOX, neither the lactate utilization genes (*lutB* and *lutC*) nor hydrogen-associated Ech complex was expressed in *Megasphaera*. The only slightly upregulated gene potentially linked to electron donor supply was encoded for aldehyde dehydrogenase (*ALDH* K00128) (log₂FC = 1.1, **Figure 6b**), which converts acetaldehyde to acetate while generating NADH (Cederbaum, 2012). Although the upstream step for acetaldehyde generation from ethanol was not detected in *Megasphaera* in our dataset (*adhE*; K13954 or K04072; **Figure S3**), previous study suggests that *Megasphaera* species can use ethanol oxidation as part of their metabolism (Wang and Yin, 2022). Its absence in our results may reflect the limited sensitivity of RNA-seq (Verwilt et al., 2023).

Another putative chain elongator *CAG-791 sp900320025* (family Lachnospiraceae) encoded a physically co-localized rBOX gene set containing *thl–hbd–crt–bcd* (**Figure S4**). Most of these genes were upregulated after THP, ranging from 1.1 to 4.7 fold except for *crt* (**Figure 6b**). Although chain elongation has not yet been reported for this exact species, other members of the Lachnospiraceae family have been shown to produce valerate and caproate through reverse β-oxidation (Fitzgerald et al., 2025). Notably, this bin showed upregulated expression of hydrogen-associated redox balancing enzyme complex Ech, together with increased expression of *ldhA* and *por*, which encode D-lactate dehydrogenase and pyruvate-ferredoxin/flavodoxin oxidoreductase, respectively (**Figure 6a and 6b**). These patterns suggested that this *CAG-791* species could rely on either hydrogen gas in the headspace or lactate produced in the feedstock to generate the reducing equivalents for rBOX. This inference is broadly consistent with the metabolic flexibility reported for a cultured member of the CAG-791 lineage, but has not been directly demonstrated for this exact species (Kaminsky et al., 2023), further experimental validation is needed to firmly connect genotype with phenotype. Overall, this analysis identified two organisms actively involved in enhanced chain elongation after THP, but because of their relatively low abundance, more evidence is needed to confirm their functions in our system.

## 4. Conclusions

In this study, applying THP to food waste increased feedstock solubility but did not increase total VFA yield. This was accompanied by reduced microbial diversity and lower acidification-related functional potential and activity at both the DNA and mRNA levels. However, the remaining community showed elevated absolute mRNA/DNA activity associated with acid production, highlighting the potential to shape microbiomes through top-down engineering control toward reduced but more functionally specialized communities for acid production. Propionate remained the dominant product regardless of THP and was consistently associated with the dominant *Prevotella* genus. However, THP reactors showed species-level enrichment of two *Prevotella* with unique functional niches. Additionally, butyrate and valerate fractions increased after THP, and two low-abundance but active chain elongators were identified. Both showed upregulated reverse β-oxidation genes and different strategies to supply the reducing equivalents required for this process.

Overall, from a process engineering perspective, this study highlights the distinctions in the utility of THP to different end objectives. While THP is useful in enhancing biogas production via anaerobic digestion, similar benefits may not be applicable while focusing on the production of VFAs. From a multi-omics perspective, this study linked molecular-level evidence to community-scale performance to identify the functional niches of individual organisms, providing valuable data and insights for future targeted microbiome engineering in complex waste-to-resource systems.

## Authorship contribution statement

MJ: Methodology, Formal analysis, Investigation, Data Curation, Writing - Original Draft, Writing - Review & Editing, Visualization

HK: Data curation, Investigation, Review & Editing, Visualization

NAD: Data curation, Investigation, Review & Editing, Visualization

KG: Data curation, Investigation, Methodology, Review & Editing

WK: Conceptualization, Review & Editing

LR: Writing - Review & Editing, Supervision

KC: Conceptualization, Methodology, Investigation, Writing - Review & Editing, Supervision, Project administration, Funding acquisition

## Acknowledgement

This study was financially supported by the Water Research Foundation project WRF 4900.

