## Supplemental Materials for "Genome-resolved multi-omics insights into top-down microbiome engineering for carbon-upcycling of thermally hydrolysed food waste"

**Contents:**

**Supplementary Tables: 7**

[Supplement Table.xlsx](https://docs.google.com/spreadsheets/d/1CsyRmrtmYqKinmzOLAjh4z1OHRdg-cJw/edit?gid=1367457421#gid=1367457421) (see the supplementary excel file for details)

**Table S1:** Three food waste batches characteristics with and without THP

**Table S2**: Assembly quality and taxonomy of the 21 assembled MAGs

**Table S3**: Presence and absence matrix of carbon metabolic functions across 21 MAGs as annotated by METABOLIC-G

**Table S4**: Customized metabolic network for individual VFA production

**Table S5**: Statistical test of the food waste characteristics between THP and non-THP

**Table S6**: Comparison of the VFA production performance between THP and non-THP

**Table S7:** Log2 fold change of DNA and mRNA involved in customized acidification networks after THP

**Supplementary Figures: 6**

**Figure S1:** 16s rRNA sequencing results of bacterial community diversity and composition in both reactors.

**Figure S2**: Comparison of mRNA, DNA, mRNA/DNA ratio and fold change of mRNA/DNA ratio in major functional categories (eggNOG) between THP and non-THP reactors

**Figure S3**: mRNA expression level of all bins within the niches for chain elongation

**Figure S4.** Gene-set organization on representative MAGs bin 18 (*Megasphaera sp000417505*) and bin 6 (*CAG-791 sp900320025*)

**Figure S5**: mRNA expression level of all bins within the niches for propionic acid production through the methylmalonyl-CoA (MMC) pathway

**Figure S6**: Comparative pangenome analysis of all assembled Prevotella MAGs

**Supplementary Methods**

### Reactor set-up and operations

- **Characterizations of food waste and measurement of reactor’s performance**
- **Calculation of performance evaluation matrix**
- **Calculation of DNA and mRNA-level comparison parameters**

#### Supplementary Figures:

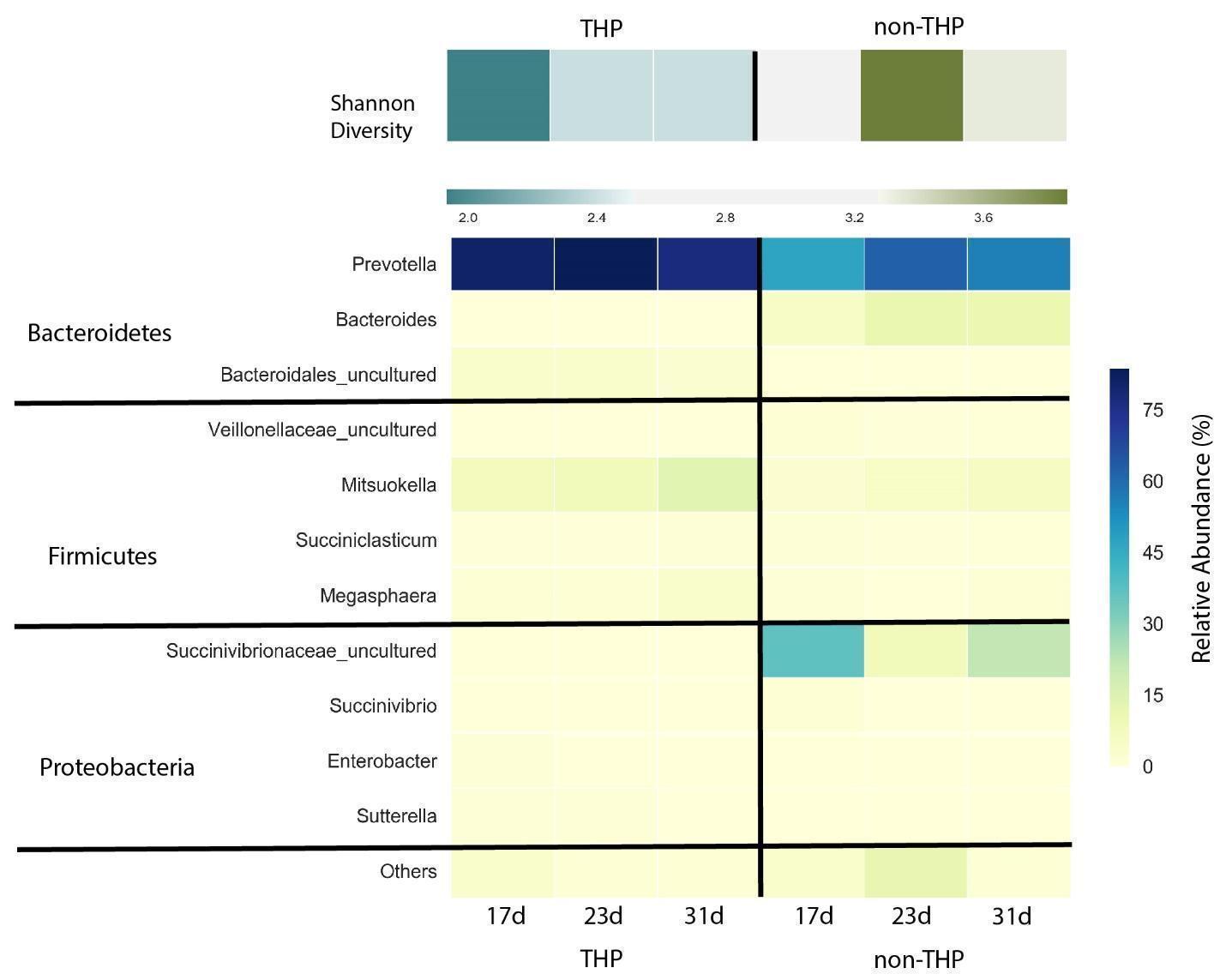

**Figure S1**. 16s rRNA gene amplicon-sequencing results of bacterial community diversity (Shannon diversity index) and composition (relative abundance on genus level) in both reactors. Relative abundances of population detected at >1% are shown, otherwise are included into others.

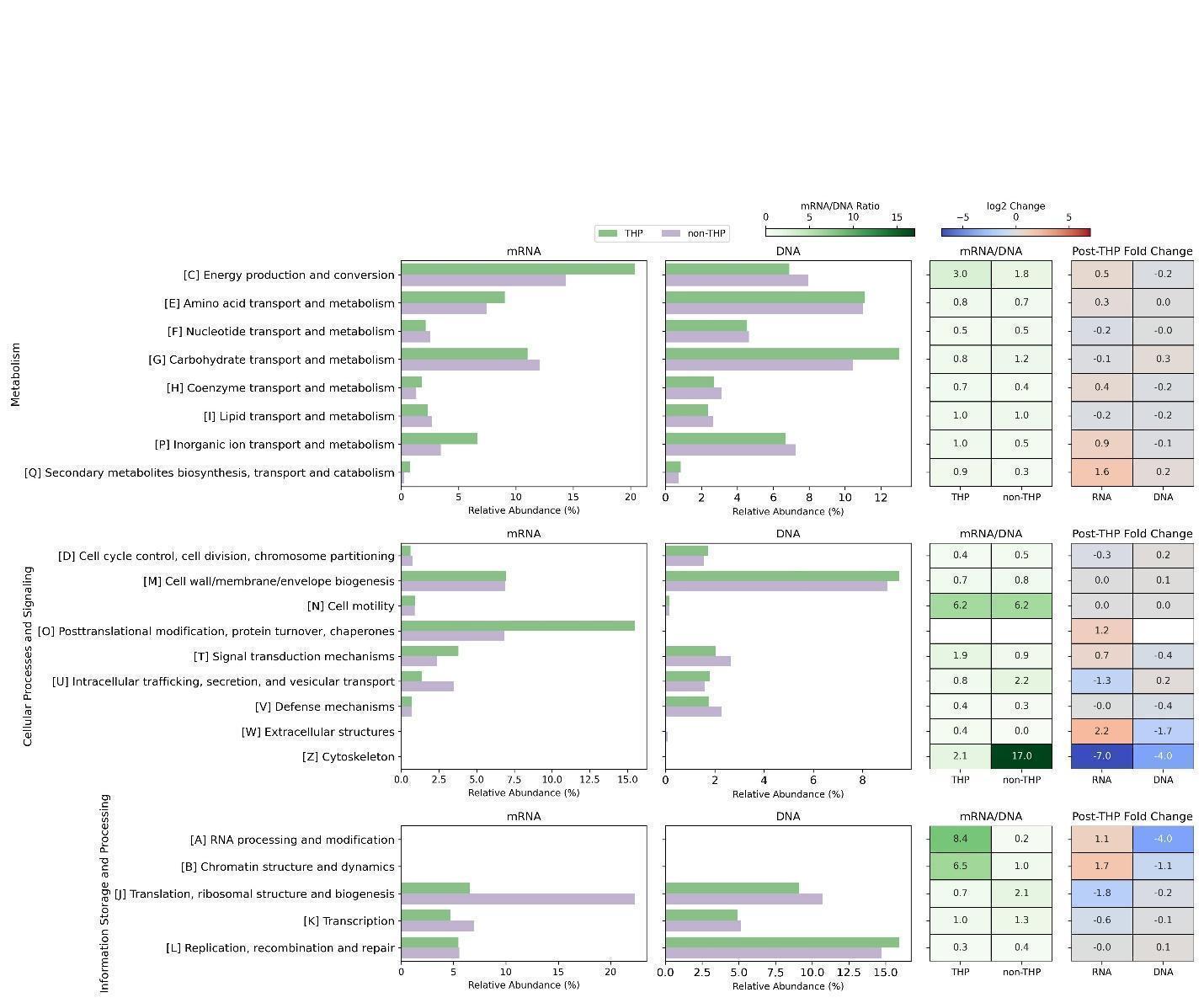

**Figure S2.** Comparison of reads-based relative abundance of mRNA, DNA, mRNA/DNA ratio and fold change of mRNA/DNA ratio in major functional categories (eggNOG) between THP and non-THP reactors.

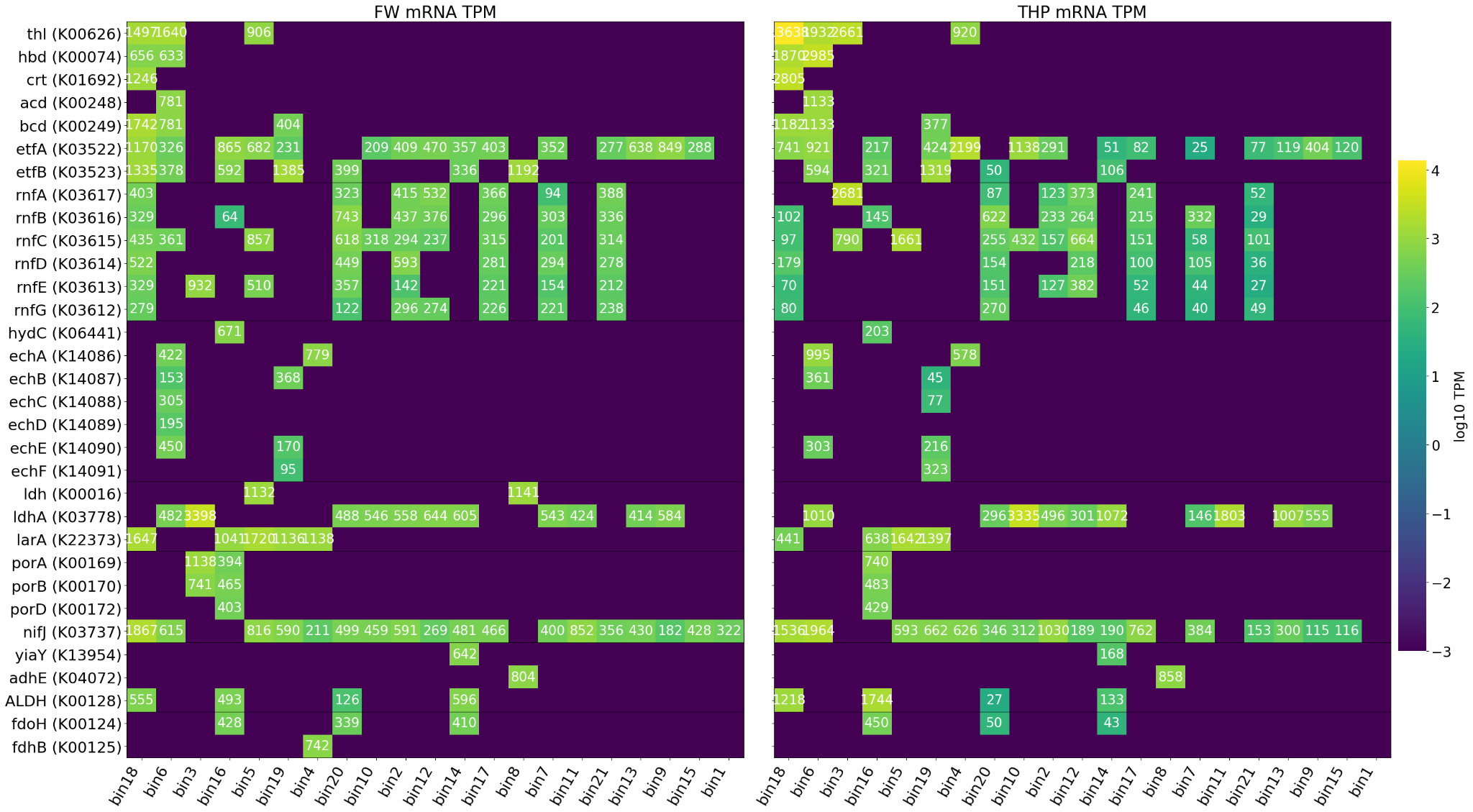
**Figure S3.** **mRNA expression levels of all bins were grouped into functional modules for chain elongation**. reverse β-oxidation (rBOX), electron-donor metabolism (e.g., lactate and ethanol oxidation to acetyl-CoA and NADH), and energy-conservation systems (Rnf and Ech complexes). Only genes expressed in at least one bin are shown (for example, Ech is not shown because none of the bins expressed genes involved in the Ech complex).

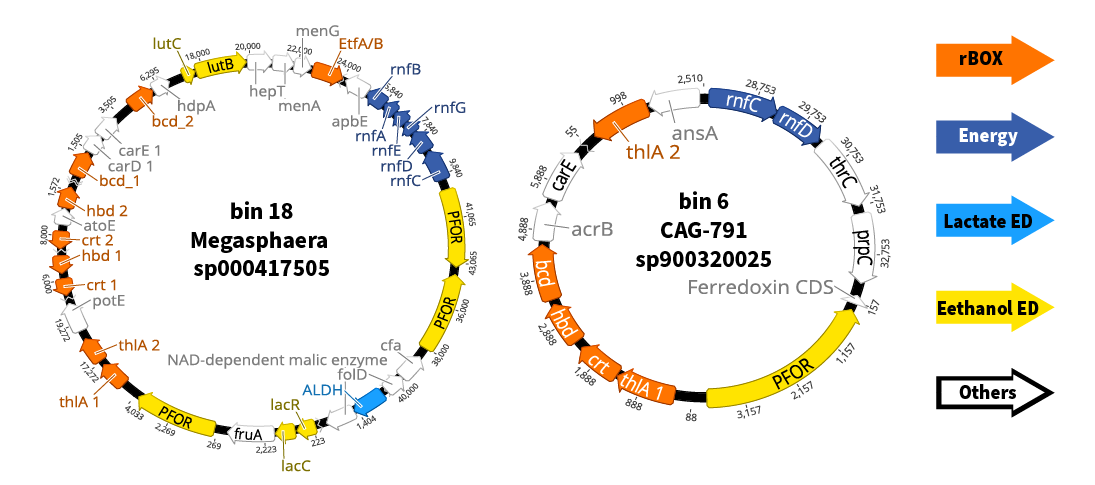

**Figure S4.** Gene-set organization on representative MAGs. Bin 18 (*Megasphaera sp000417505*) and bin 6 (*CAG-791 sp900320025*) encoded nearly complete rBOX pathways, while bin 8 (*Bifidobacterium thermacidophilum*) encoded ethanol-to-acetaldehyde conversion. Genes are colored by function: rBOX (orange), Rnf (dark blue), ethanol donor (yellow), lactate donor (blue), other (white).

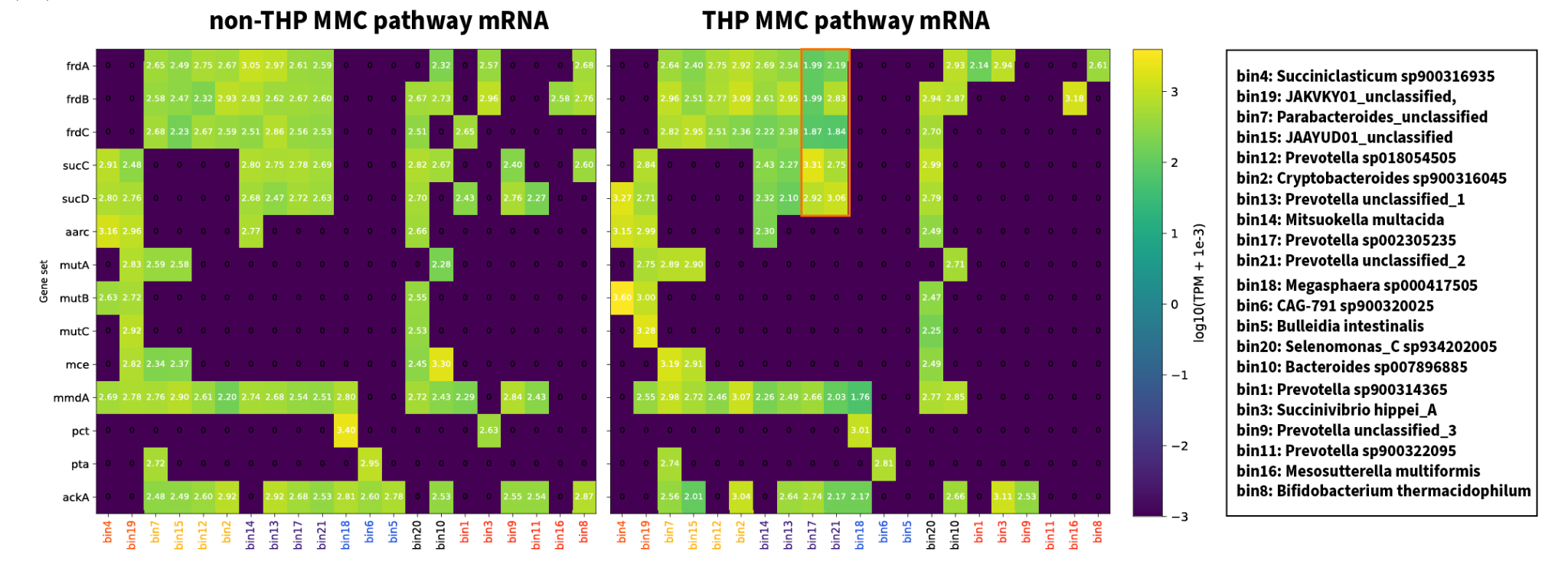

**Figure S5. mRNA expression level of all bins within the methylmalonyl-CoA (MMC) pathway of propionic acid production.** Bins are colored by similarity in the expression maps. Clustering of gene expression based on presence and absence revealed distinct groups of MAGs with incomplete or divergent propionate production capacities. For example, some MAGs lacked fumarate reductase (*frdA/B/C*) and the terminal propionyl-CoA to propionate module (bins 4 and 19), others lacked *sucC/sucD* (succinyl-CoA synthetase) (bins 7, 15, 12, and 2) or *mutA/B/C* and *mce*, disrupting the succinyl-CoA to propionyl-CoA node (bins 14, 13, 17, and 21), while a subset contained only downstream modules converting propionyl-CoA to propionate (bins 18, 6, and 5). Some MAGs showed condition-specific variation, such as bin 10, which actively expressed a nearly complete pathway under non-THP but lost most modules under THP.

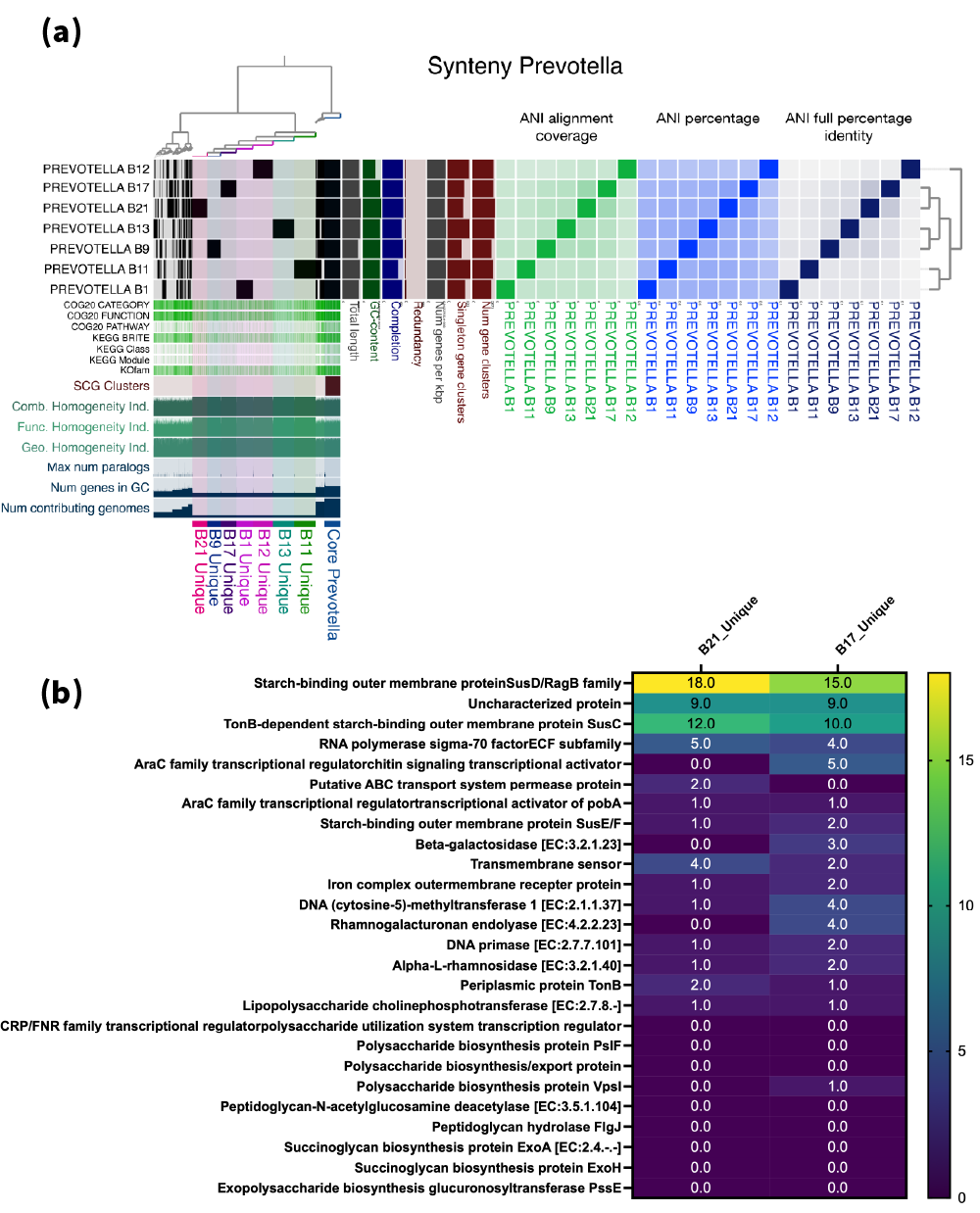

**Figure S6. Comparative pangenome analysis of all assembled Prevotella MAGs. (a)** Pangenome synteny map of *Prevotella* MAGs showing unique gene sets in each bins (outer rings), annotated by COG20, KEGG pathway, BRITE, and KOfam categories. Bin 17 (*Prevotella sp002305235*) and bin 21 (*Prevotella unclassified_2*) cluster closely, reflecting shared genomic content. The left panel shows synteny-based hierarchical clustering of seven *Prevotella* bins using gene order conservation and homologous cluster content. Colored blocks summarize genomic features including COG/KEGG category distributions, ANI alignment coverage, ANI identity, gene family redundancy, and homologous cluster sharing among bins. **(b)** All Unique orthology groups annotated by KOfam in bins 17 and 21.

**Supplementary Methods:**

### Reactor set-up and operations

Two 6-L semi-continuous reactors were operated at 37.0 ± 2.0$℃$ for 38 days, fed separately with THP- and non-THP treated food waste. The pH was maintained at 6.20 ± 0.05. An inline probe connected to a transmitter (Mettler Toledo M400) continuously monitored pH and automatically triggered a dosing pump to add 1M NaHCO₃ when pH fell below the set point. The same probe system also monitored ORP and temperature (Mettler Toledo pH/ORP probe). The organic loading rate (OLR) was targeted at 6.25g total COD/L/d, and the hydraulic retention time (HRT, equivalent to the solids retention time SRT in our reactors), was maintained at 2 days to limit methanogen growth. This semi-continuous reactor operation and the targeted OLR and HRT were achieved by setting up a timer and automatic pumping system in which effluent was first withdrawn. Specifically, within 5 min, 3 L of well-mixed solution was pumped out from each 6-L bioreactor, followed by influent pumping with 3 L of thawed influent food waste. The pumping speeds were optimized to balance minimal feeding time against the risk of tube clogging and pumping failures.

- **Characterizations of food waste and measurement of reactor’s performance**

The influent feedstock was characterized by their chemical oxygen demand (COD), total solids (TS), volatile solids (VS), and the organic compositions (carbohydrate, proteins, lipids). The soluble compounds were quantified after removing the particles using 0.22-micrometer filters. COD was quantified using COD Digestion Vials (high range, Hach). Total solids (TS) and volatile solids (VS) were measured following American Public Health Association Standard Methods for Examination of Water and Wastewater (2012). Carbohydrate, protein, and total lipids were quantified using the anthrone method (Morris, 1948), modified Lowry method (Thermo Fisher Scientific, IL, USA), and Folch method (Folch et al., 1957), respectively. Total ammonia nitrogen (TAN) concentration was measured using an ammonia electrode (Thermo Scientific, Sunnyvale, CA), and the total carboxylic acid (tCA) was evaluated using the Hach company esterification methods 8196.

The measurements of reactor’s performance including the soluble COD (sCOD), VFAs and produced methane gas in the effluent. Soluble COD (sCOD) was measured following the same method for influent. VFAs were measured by [Ion chromatography](https://www.sciencedirect.com/topics/biochemistry-genetics-and-molecular-biology/ion-chromatography) (Dionex 2000 system equipped with an AS11 column). The analyzed VFAs included acetic acid, propionic acid, butyric acid, and valeric acid. Methane concentrations in the bioreactor headspace were analyzed using [gas chromatography](https://www.sciencedirect.com/topics/agricultural-and-biological-sciences/gas-chromatography) (Model 8610C, SRI instruments) with a ShinCarbon ST column and [thermal conductivity detector](https://www.sciencedirect.com/topics/biochemistry-genetics-and-molecular-biology/thermal-conductivity-detector) (TCD).

- **Calculation of performance evaluation matrix**

$$VFA conversion efficiency \left( \% \right)=\frac{Effluent COD\left( VFA \right)-Influent COD \left( VFA \right)}{Influent tCOD}\times100\%$$

$$\boldsymbol{Methane yield (\%) =}\frac{\boldsymbol{Total COD}\left( \boldsymbol{C}\boldsymbol{H}_{\boldsymbol{4}} \right)}{\boldsymbol{Influent tCOD}}\boldsymbol{\times100\%}$$

$$\boldsymbol{VFA accumulation level (\%) =}\frac{\boldsymbol{Total COD of all measured VFAs}}{\boldsymbol{Influent tCOD}}\boldsymbol{\times100\%}$$

$$\boldsymbol{Individual acid proportions (\%) =}\frac{\boldsymbol{individual acid concentration \times COD equivalent conversion factor}}{\boldsymbol{Sum of four acids COD equivalent}}\boldsymbol{\times100\%}$$

The COD conversion factors were used as followed: 4g COD/g CH4, 1.07 g COD/g acetate, 1.51g COD /g propionate, 1.82g COD/ g butyrate, and 2.04 g COD/g valerate. All values were calculated after both reactors reached a relatively stable performance (the effluent VFA concentration had less than 5% relative standard deviation (RSD)).

- **Calculation of DNA and mRNA-level comparison parameters**

$\boldsymbol{Global function DNA reads relative abundance (\%) =}\frac{\boldsymbol{number of mapped DNA reads per global functional classification}}{\boldsymbol{total number of DNA reads per sample}}$

$\boldsymbol{Global function mRNA reads relative abundance (\%) =}\frac{\boldsymbol{number of mapped mRNA reads per global functional classification}}{\boldsymbol{total number of mRNA reads per sample}}$

$\boldsymbol{DNA TPM=}\frac{\boldsymbol{number of mapped DNA reads per gene \times}\boldsymbol{10}^{\boldsymbol{6}}}{\frac{\boldsymbol{gene length}}{\boldsymbol{1kb}}\boldsymbol{\times total number of DNA reads per sample}}$

$\boldsymbol{mRNA TPM=}\frac{\boldsymbol{number of mapped mRNA reads per gene \times}\boldsymbol{10}^{\boldsymbol{6}}}{\frac{\boldsymbol{gene length}}{\boldsymbol{1kb}}\boldsymbol{\times total number of mRNA reads per sample}}$

$\boldsymbol{Per-genome transcriptional levels =}\frac{\boldsymbol{DNA TPM}}{\boldsymbol{mRNA TPM}}$
